# An endocannabinoid-nitric oxide signaling switch triggers reciprocal inhibitory-excitatory plasticity at dopamine neuron inputs following prenatal cannabinoid exposure

**DOI:** 10.64898/2026.09.09.750396

**Authors:** Valeria Serra, Federico Brandalise, Vivien Miczán, István Katona, Miriam Melis

## Abstract

Dynamic regulation of midbrain dopamine neuron activity is necessary for diverse processes including motivation, novelty detection, reinforcement learning, and cognitive flexibility. By setting the strength of synaptic inputs to dopaminergic neurons, endocannabinoid signaling is essential for regulating dopaminergic activity. Prenatal exposure to Δ9-tetrahydrocannabinol (THC), the main psychoactive substance in cannabis, is known to induce abnormal dopaminergic activity and increased susceptibility to psychopathology. However, how prenatal cannabinoid exposure (PCE) affects endocannabinoid-mediated synaptic plasticity of dopamine neurons remains largely unknown. Here, we use a rat model of PCE to directly determine this. We found that endocannabinoid-mediated synaptic plasticity at excitatory synapses on dopamine neurons of the ventral tegmental area (VTA) was absent in PCE male rat offspring, where the presynaptic nanoscale architecture of excitatory afferents onto VTA dopamine neurons was reorganized to impair the control of type-1 cannabinoid receptors on glutamate release. We demonstrate that, in response to postsynaptic depolarization, PCE dopamine cells switch to nitric oxide (NO) rather than endocannabinoid signaling to induce opposing forms of synaptic plasticity at inhibitory and excitatory inputs. We disclosed a NO-dependent long-term potentiation of GABA*_A_*-receptor-mediated synaptic transmission, and a long-term depression of glutamatergic synapses requiring presynaptic activation of GABA*_B_*-receptors. These PCE-induced reciprocal forms of synaptic plasticity reshape the balance of excitatory and inhibitory control over dopamine neurons, potentially contributing to increased vulnerability to psychiatric disorders.

**Significance Statement:** Endocannabinoids (eCBs) in the midbrain regulate dopamine cell activity and plasticity to guide behavior. eCBs retrogradely activate presynaptic type-1 cannabinoid receptors to depress synaptic transmission. Here, we identify metaplastic changes driven by prenatal cannabinoid exposure (PCE) on eCB signaling at excitatory afferents on dopamine neurons. These include a switch in retrograde signaling favoring de novo synthesis of nitric oxide (NO) to replace the actions of eCBs, which are compromised at excitatory inputs. NO induces novel forms of long-term plasticity at excitatory and inhibitory inputs on male dopamine neurons. These forms of metaplasticity may be recruited in the VTA in response to PCE-induced circuit remodeling, and the resulting synaptic adaptations contribute to abnormal dopamine cell activity.

## 1. Introduction

Cannabis use during pregnancy has risen sharply coinciding with its increasing normalization and social acceptance (1–4). Its expanding legal availability is leading to the common public misconception that it is a safe natural remedy to be used even during vulnerable periods such as pregnancy (5–8). However, type-1 cannabinoid (CB_1_) receptors are expressed in the embryonic brain where their activation by Δ9-tetrahydrocannabinol (THC), the main psychoactive ingredient in cannabis, can alter axonal growth and guidance (9, 10). Accordingly, prenatal cannabis exposure (PCE) is a predictive risk factor for the development of psychopathologies in the offspring, including psychotic-like experiences (2, 11–20). In children, PCE also enhances the risk for the development of overt psychotic disorder (12, 13, 20) with a hallucinatory-like profile (20). Of note, hallucination-like percepts are tied to an increased dopaminergic state (21, 22). In psychotic disorders, one of the most well-established neurophysiological abnormalities is detrimental sensorimotor gating functions (23). Notably, in a rat model of PCE, we have previously identified a male-specific latent vulnerability that manifests as impaired sensorimotor gating functions following a single exposure to THC (24) or acute unavoidable stressors (25, 26). Such deficits are tied to an increased dopamine neuronal activity within the ventral tegmental area (VTA) (24, 26, 27).

Retrograde signaling, finely regulating synaptic strength within the VTA, might contribute to adjusting the weight of plasticity via activation of presynaptic type-1 cannabinoid (CB_1_) receptors. Hence, early alterations to endocannabinoid signaling in the dopaminergic system may contribute to vulnerability in PCE progeny. Here, we tested the hypothesis that PCE triggers metaplastic changes in retrograde endocannabinoid signaling at excitatory synapses in the VTA. By using electrophysiological, microscopic and imaging techniques, we found that a short-term form of plasticity mediated by endocannabinoids in the VTA and expressed by dopamine neurons at glutamatergic synapses was absent in PCE males. Glutamatergic synapses also show an impairment of CB_1_ signaling. Instead, upon depolarization, PCE male dopamine cells release nitric oxide (NO) to drive a novel form of long-term depression (LTD) at these afferents that requires activation of presynaptic GABAb receptors. Concurrently, PCE male dopamine cells express a NO-dependent form of long-term potentiation (LTP) at inhibitory synapses. Such PCE-induced metaplastic mechanisms, involving the recruitment of a different retrograde messenger when CB1 receptor function is compromised, might be key not only in offsetting the excitation–inhibition balance but also in understanding the underlying neurobiological mechanisms that lead to the behavioral consequences of maternal cannabis use in offspring.

## 2. Materials and Methods

### 2.1 Animals

All experimental procedures were performed in accordance with the European legislation EU Directive 2010/63 and the Animal Ethics Committees of the University of Cagliari and by the Italian Ministry of Health (authorization number: 256/2020). We made all efforts to minimize pain and suffering and to reduce the number of animals used. Primiparous female Sprague Dawley rats (Envigo) were used as mothers and single housed during pregnancy. THC or vehicle was administered subcutaneously (2 mg/Kg, 1 mL/Kg, s.c. once per day) from gestational day (GD) 5 to GD 20(24). Offspring were weaned at postnatal day (PND) 21 and were housed in a standard condition of temperature (21 ± 1 °C) and humidity (60 %) under a normal 12 h light–dark cycle with ad libitum access to water and food until the experimental day (PND 15-28). To control for litters effects, we did not use more than two offsprings from each litter for the same experiment.

### 2.2 Electrophysiological recordings

The preparation of the lateral posterior VTA slices was performed as previously described(24). Briefly, horizontal midbrain slices (250 µm) containing the VTA was obtained from male and female offspring anesthetized with isoflurane. The tissue was sliced with a vibratome (Leica) in ice-cold low Ca2+ solution containing the following (in mM): 126 NaCl, 1.6 KCl, 1.2 NaH2PO4, 1.2 MgCl2, 0.625 CaCl2, 18 NaHCO3 and 11 glucose (304-306 mOsm). Immediately after cutting, slices were transferred to a holding chamber (37 °C) with artificial cerebrospinal fluid (ACSF) saturated with 95% O2 and 5% CO2 containing the following (in mM): 126 NaCl, 1.6 KCl, 1.2 NaH2PO4, 1.2 MgCl2, 2.4 CaCl2, 18 NaHCO3 and 11 glucose (304-306 mOsm). Slices were allowed to recover for at least 1 h before being placed, as hemislices, in the recording chamber and superfused with ACSF (36-37 °C) saturated with 95% O_2_ and 5% CO_2_. Cells were visualized using an upright microscope with infrared illumination (Axioskop FS 2 plus, Zeiss), and whole-cell patch-clamp recordings were made using an Axopatch 200B amplifier (Molecular Devices). Voltage-clamp recordings of evoked EPSCs were made with electrodes filled with a solution containing the following (in mM): 117 caesium methanesulfonic acid, 20 HEPES, 0.4 EGTA, 2.8 NaCl, 5 TEA-Cl, 0.1 mM spermine, 2.5 Mg2ATP and 0.25 Mg2GTP, pH 7.2-7.4, 275-285 mOsm. Voltage-clamp recordings of evoked IPSCs recordings were made with electrodes filled with a solution containing the following (in mM): 144 KCl, 10 HEPES buffer, 3.45 BAPTA, 1 CaCl2, 2.5 Mg2ATP and 0.25 Mg2GTP, pH7.2-7.4, 275-285 mOsm. Experiments were begun only after series resistance had stabilized (typically 10-30 MO). Series and input resistance were monitored continuously on-line with a 5 mV depolarizing step (25 ms). Data were filtered at 2 kHz, digitized at 10 kHz, and collected on-line with acquisition software (pClamp 10; Molecular Devices). Dopamine neurons from the lateral portion of the posterior VTA were identified by the following criteria: cell morphology and anatomical location to the medial terminal nucleus of the accessory optic tract, the presence of a large hyperpolarization-activated current (> 100 pA) assayed immediately after break-in using 13 incremental 10 mV hyperpolarizing steps from a holding potential of −70 mV. A bipolar, stainless steel stimulating electrode (FHC) was placed −100-200 μm rostral to the recording electrode and was used to stimulate at a frequency of 0.1 Hz. Paired stimuli were given with an interstimulus interval of 50 ms, and the ratio between the second and the first postsynaptic currents (PSC2/PSC1) was calculated and averaged for a 5-min baseline. The depolarizing pulse used to evoke depolarization-induced suppression of excitation (DSE) was a 5 s step to +40 mV from holding potential(28). This protocol was chosen on evidence of an endocannabinoid tone when DA cells are held at +40 mV(28). The magnitude of DSE was measured as a percentage of the mean amplitude of consecutive PSCs after depolarization (acquired between 5 and 15 s after the end of the pulse) relative to that of 5 min of baseline mean amplitude acquired before depolarization. The potentiation/depression of evoked PSCs induced by depolarization were measured as a percentage of the mean amplitude of consecutive PSCs recorded for 30 min by the average 10 min of baseline immediately prior to the depolarization step.

### 2.3 Nitric oxide imaging using DAF-FM diacetate

Nitric oxide (NO) production was assessed in acute coronal midbrain slices containing the ventral tegmental area (VTA) using the NO-sensitive fluorescent probe DAF-FM diacetate. Coronal slices (250–300 µm) were prepared and maintained in continuously oxygenated artificial cerebrospinal fluid (aCSF; 95% O₂ and 5% CO₂). Slices were incubated with DAF-FM diacetate (10 µM in oxygenated aCSF) for 30 min in the dark to allow intracellular loading and de-esterification. After loading, slices were transferred to the imaging chamber and continuously superfused with oxygenated aCSF throughout the experiment. Fluorescence imaging was performed on an epifluorescence microscope equipped with a Teledyne Photometrics (Tucson, AZ, USA) Prime BSI Express sCMOS camera. Excitation was provided by an LED light source and fluorescence was collected through a standard FITC filter set. Images were acquired using a 40× water-immersion objective (Olympus LUMPlanFLN 40×/0.80 W) under identical acquisition settings across experimental conditions (illumination intensity, camera readout configuration, binning, exposure time, and field of view). Time-lapse recordings were acquired at 4 Hz (250-ms inter-frame interval) for 60 consecutive time points (total duration ≈ 15 sec), comprising a pre-stimulus baseline, imaging during a brief depolarizing stimulus, and a post-stimulus period. To minimize photobleaching and phototoxicity while preserving temporal alignment with the stimulus, exposure time was set to 50 ms and LED intensity was kept at the lowest level yielding adequate signal-to-noise ratio and maintained constant across groups. Regions of interest (ROIs) were defined as circular areas centered on the soma of the recorded cells. To minimize variability due to dye loading and optical factors, analyses focused on within-ROI stimulus-evoked fluorescence changes. Images were background-subtracted and responses were quantified as ΔF/F₀, where F₀ was defined as the mean fluorescence during the pre-stimulus baseline for each ROI. Baseline fluorescence values were not used to infer differences in basal NO levels between groups. Recordings displaying visible movement, focus drift, uneven illumination, or unstable baseline fluorescence (e.g., progressive monotonic decay consistent with bleaching) were excluded according to pre-defined criteria.

### 2.4 Immunostaining and confocal image analysis

Immunostaining and imaging were performed according to (58). Briefly, rats were transcardially perfused with 4% (m/v) paraformaldehyde (PFA) or brain slices were immersion-fixed in 4% PFA overnight, and sectioned to 20 μm in phosphate buffer (PB) using a Leica VT-1000S Vibratome for STORM imaging and 50 μm for confocal analysis. Immunostaining was performed in a free-floating manner. PB and 0.05 M Tris-buffered saline (TBS, pH 7.4) washes were followed by a blocking and permeabilizing step in 5% (v/v) normal donkey serum (Sigma) and 0.3% (v/v) Triton X-100 (Sigma) in TBS for 45 min, then slices were incubated in primary antibodies ovenight (see Supplementary Table X) in TBS. Sections were then washed in TBS and incubated with the appropriate secondary antibodies (see Supplementary Table X) supplemented with 4,6-diamidino-2-phenylindole (DAPI; 1:1,000), if needed, then washed in TBS and PB.

Sections were mounted in VectaShield (Vector Laboratories) or Prolong Diamond Antifade Mounting Medium (Invitrogen) for confocal imaging on a NIKON A1R microscope (voxel size: 0.5 × 0.5 × 0.15 um). A region of interest of 300×300 um was defined in the VTA and a deep-learning-based single-cell segmentation algorithm (Generic cytoplasm segmentation v1) was applied on the images with the Biological Image Analysis Software (Single-Cell Technologies Ltd.). Then, after feature extraction, a machine learning model was trained to distinguish TH+/nNOS+, TH+/nNOS− and TH−/nNOS+ cells along with a fourth class comprising cell debris and fluorescence-related artifacts which was excluded from further analysis. Segmentation and classification results were checked manually to ensure quality. Cell density, along with fluorescent intensity levels of the corresponding channels were automatically calculated using the segmentation masks and aggregated over the regions of interests for the corresponding classes.

### 2.5 Correlated confocal and STORM imaging processing

For STORM imaging, sections were post-fixed in 4% PFA for 10 min and washed in PB, then mounted and dried on acetone-cleaned no. 1.5 borosilicate coverslips.

Before imaging, STORM imaging medium (containing 0.1 M mercaptoethylamine, 5% (m/v) glucose, 1 mg ml^−1^ glucose oxidase and catalase 2.5 μl ml^−1^ of aqueous solution from Sigma, approximately 1,500 U ml^−1^ final concentration in Dulbecco’s PBS from Sigma was freshly prepared as prevously described (59). Coverslips were sealed with nail polish.

Imaging was performed 10 min – 180 min after sample preparation using a Nikon Ti-E inverted microscope equipped with a Nikon N-STORM system, CFI Apo TIRF ×100 objective (1.49 numerical aperture), a Nikon C2 confocal scan head and an Andor iXon Ultra 897 EMCCD (with a cylindrical lens for astigmatic 3D-STORM imaging (60). Imaging process was controlled by Nikon NIS-Elements AR software with the N-STORM module using a 300-mW laser (VFL-P-300-647, MPB Communications) fiber-coupled to the laser board. vGluT1-positive axon terminals impinging on TH−positive cell bodies and dendrites were selected using the live EMCCD image with a 488-nm illumination, then a three-channel confocal stack (512 × 512 × 15 pixels, 78×78×150nm resolution) was then collected using 488-nm, 561-nm and 647-nm excitations. After brief bleaching, direct STORM imaging was performed with 10,000 cycles of 30 ms of exposure, with continuous low-power activator laser (405 nm) and maximal power reporter laser (647 nm) using a STORM filter cube (Nikon) and the EMCCD camera. Huygens software (SVI) was used to deconvolve confocal stacks with 100 iterations of the classic maximum likelihood estimation algorithm and STORM image processing was performed using the N-STORM module of the NIS-Elements AR with a peak detection threshold of 1,000 gray levels. Correlated confocal and STORM image analysis was performed using the VividSTORM software (58). The two imaging modalities were manually aligned based on the correlated confocal and STORM channels. The borders of the axon terminals and active zones were delineated on the confocal images using the Morphological Active Contour Without Edges (MACWE) algorithm. Number of STORM localization points (LPs) was calculated based on the ROIs and were normalized to the overall density of LPs per image. Density of bassoon staining in the active zone was calculated by dividing the bassoon NLPs and the active zone size. Figures were prepared using Adobe Photoshop (24.2.1.) and Adobe Illustrator (29.8.1). All images were modified in the same way for all treatment groups during preparation of the figures to ensure equal comparison.

### 2.6 Drugs

THC resin was purchased from THC PHARM GmbH (Frankfurt, Germany) and dissolved in ethanol at 20 % final concentration. Then, THC was suspended in a vehicle (VEH) solution containing 1–2% Tween® 80 and diluted with sterile saline (0.9 % NaCl). WIN 55,212-2, HU210, ODQ and CGP35348 were purchased from Sigma-Aldrich. AM281, L-NAME, NPLA and DAF-FM diacetate were purchased from Tocris Bioscence. All drugs were dissolved in dimethyl sulfoxide (DMSO) when it was needed. Particularly, WIN, 55,212-2 and AM281 were dissolved in DMSO. The final concentration of DMSO was <0.01%.

### 2.7 Statistical analysis

All the numerical data are given as mean ± SEM. Statistical analysis was performed using GraphPad Prism (version 8). Statistical outliers were identified using Grubb’s test (α = 0.05) and excluded from analyses. For STORM imaging, the mean values of each animal were used in the statistical analyses, differences between the groups were determined using Mann–Whitney U-tests. Data always met the assumptions of the applied statistical probe. Electrophysiological data were analyzed using two-way ANOVA or two-way ANOVA for repeated measures (treatment × time) or Student’s t-test when appropriate, followed by Sidak’s, or Tukey’s post hoc test when the interaction between factors was revealed. The significant threshold value was set at 0.05.

## 3. Results

### 3.1 In utero THC exposure disrupts CB_1_-dependent signaling in male VTA

Endocannabinoid signaling via CB_1_ receptor activation leads to a presynaptic inhibition of glutamatergic transmission in VTA dopaminergic neurons (28, 29). To test if PCE alters endocannabinoid signaling at these synapses, we elicited a CB_1_ receptor-dependent form of short-term synaptic plasticity, termed depolarization-induced suppression of excitation (DSE) (Fig. 1A) (28, 30). We recorded AMPA-receptor-mediated excitatory post-synaptic currents (EPSCs) from control (CTRL) male dopamine cells that displayed a DSE, which was similar in magnitude to naïve males (28, 30) (Fig. 1B-D), while PCE male dopamine neurons exhibited a larger DSE (~50% vs ~20% in CTRL; Fig.1B-D). An increased paired-pulse ratio (PPR) accompanied DSE in both groups (Fig 1E) suggestive of a presynaptic locus of action. In CTRL male dopamine cells, DSE required activation of CB_1_ receptors, since it was prevented by bath application of the CB_1_ receptor antagonist AM281 (0,5 μM; Fig. 1F), which also abolished PPR changes (Fig. 1G). In contrast, unexpectedly, PCE male dopamine cells expressed DSE (~50%) even in the presence of AM281 (Fig. 1F), an effect accompanied by an increased PPR (Fig. 1G). AM281 increased the amplitude of AMPA EPSCs exclusively in PCE male dopamine cells (Fig. 1H), suggestive of an endogenous tone at these afferents. Notably, upon depolarization, the amplitude of AMPA EPSCs recorded from PCE neurons did not return to baseline levels within seconds like in CTRL cells (Fig. 1I) and naïve animals (28), but it was reduced for at least 30 min, thus featuring a novel form of LTD (Fig. 1J). Because this newly recruited LTD is CB_1_ receptor-independent, it may substitute in PCE male dopamine neurons for the conventional LTD induced by pairing mild postsynaptic depolarization (−40 mV) with low-frequency stimulation (LFS; 1 Hz, 6 min; Fig. S1A–C), which is abolished after PCE (23) but normally requires CB_1_ receptor activation in naïve rats, as demonstrated by its blockade with AM281 (Fig. S1B) and occlusion by bath-applied WIN 55,212-2 (WIN; Fig. S1C).

**Figure 1.**
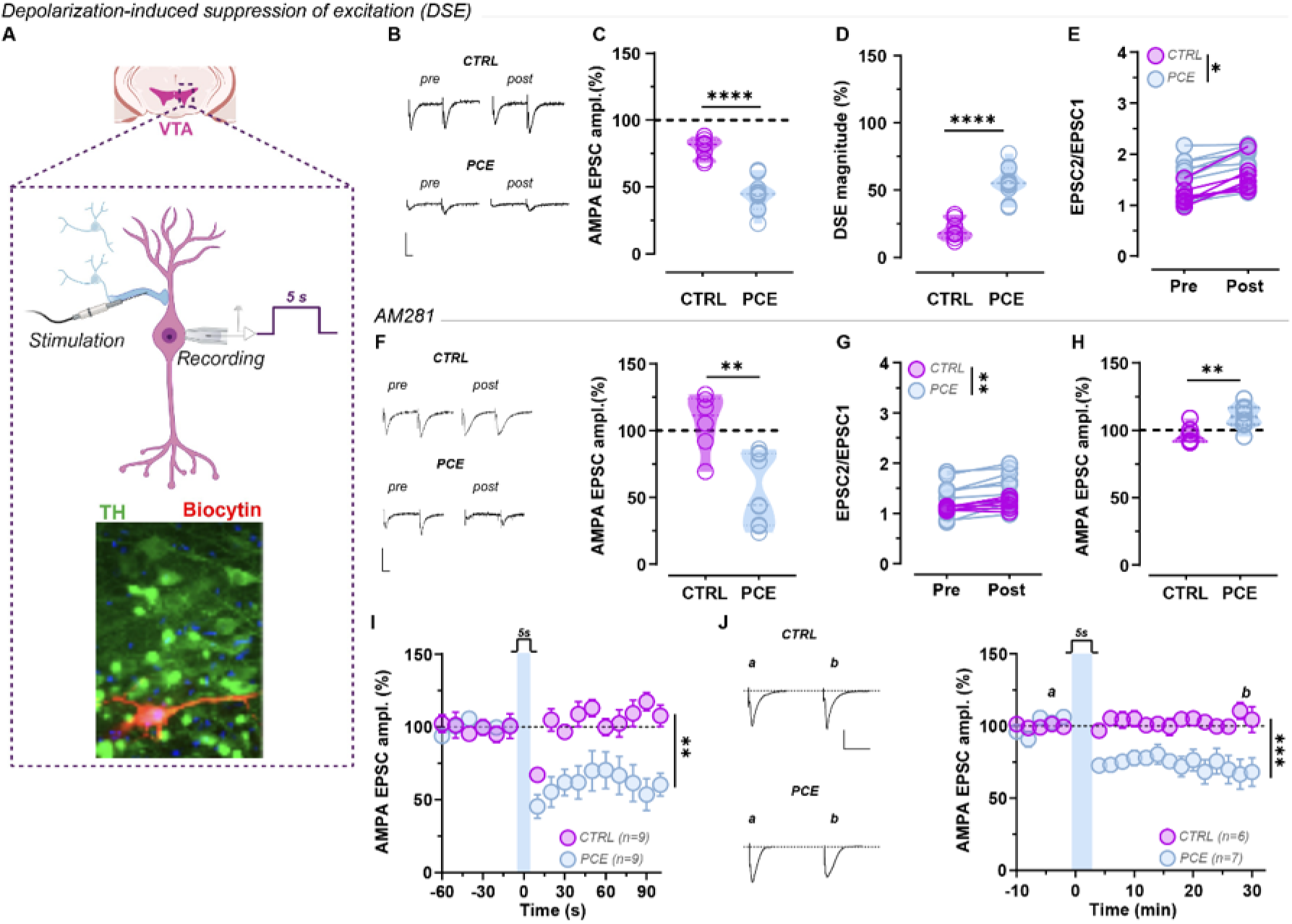
Prenatal THC decreases CB1 control of efficacy of excitatory inputs in male progeny. A) Schematic representation of experimental design showing a midbrain section with VTA in which a dopaminergic cell (DA) is recorded in response to 5 second (s) of depolarization. Inset show immunohistochemical images of co-localization of TH+ cell filled with biocytin. B) Panel shows representative traces of paired EPSCs recorded pre and post 5s depolarization from male CTRL and PCE dopamine neurons. Calibration bar, 10ms, 100pA. C, D) Graphs show the effect of 5 s depolarization on AMPA EPSC amplitude (C) and magnitude of DSE (D) in CTRL and PCE male dopamine cells (C,D; Unpaired *t*-test: t=6.602, df=15, *P* < 0.0001; *N_CTRL_* = 7 *and N_PCE_* = 10). EPSC amplitude and DSE magnitude were normalized to the averaged value (dotted line) before depolarization. Graphs show violin plots (including the median values, and lower and upper quartiles) with each circle representing a single value. E) In male dopamine cells, DSE differently increases the paired-pulse ratio (EPSC2/EPSC1) of AMPA EPSCs between PCE and CTRL (two-way ANOVA: Pre vs post’s effect: *F*_(1,14)_=45.32, *P* < 0.0001; Treatment’s effect: *F*_(1,14)_=5.828, *P* = 0.03; Interaction between factors: *F*_(1,14)_=1.131, *P* = 0.305; *N_CTRL_* = 7 and *N_PCE_* = 9). Graph plots the paired-pulse ratio for each of the experiments in C and D before (pre) and 5 s after depolarization (post). F) Graph shows that CB1R antagonist AM281 (500 nM) prevents DSE expression in dopamine cells of CTRL males, but not in PCE males (Unpaired *t*-test: t=3.843, df=13, *P* = 0.002; *N_CTRL_* = 6 and *N_PC__E_* = 9). Inset shows representative traces of EPSCs recorded pre and post DSE. Calibration bar, 10 ms, 100 pA. Graphs show violin plots (including the median values, and lower and upper quartiles) with each circle representing a single value. G) The graph plots the paired-pulse ratio for each of the experiments in F before (pre) and after DSE (post) in male dopamine neurons (Two-way ANOVA: pre vs post’s effect: *F*_(1,27)_=65.48, *P* < 0.0001; treatment’s effect: *F*_(3,27)_=5.33, *P* = 0.005; interaction between factors: *F*_(3,27)_=3.60, *P* = 0.026; Sidak’s multiple comparison test: PCE pre vs post: *P* = 0.0025; *N_CTRL_* = 6; *N_PCE_* = 9). H) Graph shows the effect of AM281 on AMPA EPSC amplitude in PCE and CTRL DA cells (Unpaired *t*-test with Welch’s correction: : t=3.103, df=11.94, *P* = 0.0092; *N_CTRL_* = 6 *and N_PCE_* = 8). I) Time course of DSE in male dopamine cells (two-way ANOVA: treatment’s effect: *F*_(1,16)_= 13.21, *P* = 0.0022; time’s effect: *F*_(5.13,82.09)_= 6.246, *P* < 0.0001; interaction between factors: *F*_(15,240)_= 6.489, *P* < 0.0001). Each point represents the average of the mean EPSCs for the 10 sec-bin (±SEM) obtained from different cells. Number in brackets indicates the number of cells. J) *Left*, traces show evoked AMPAR EPSCs obtained before (a) and after 30 min (b) of 5s depolarization. Calibration bar, 10ms, 100pA. *Right,* in PCE male dopamine cells (*N* = 7), 5s depolarization induces a long-term depression of AMPA EPSCs (two-way ANOVA: treatment’s effect: *F*_(1,11)_ = 25.61, *P* = 0.0004; time’s effect: *F*_(2.283,25.11)_= 3.085, *P* = 0.05; interaction between factors: *F*_(18,198)_= 4.769, *P* < 0.0001). Each point represents the average of the mean EPSCs for 2 min-bin (±SEM) obtained from different cells. Number in brackets indicates the number of cells. *\*P* < 0.05; *\*\*P* < 0.01; *\*\*\*P* < 0.001; *\*\*\*\*P* < 0.0001.

Previously, we found that PCE modifies the nanoscale organization and signaling of presynaptic CB_1_ receptors at inhibitory synapses on male dopamine neurons (24). To determine if and how PCE affects CB_1_ receptor function at glutamatergic synapses, the effects of increasing concentrations of two CB_1_ receptor agonists, WIN and HU210 (HU), were measured on AMPA EPSCs recorded from dopamine cells. Both drugs produced a dose-dependent reduction of AMPA EPSCs in CTRL dopamine cells (Fig. 2A) that was similar to that observed in naïve rats (28). However, this effect was markedly decreased on AMPA EPSCs recorded from PCE male dopamine cells (Fig. 2A), thus indicating less effective CB_1_ receptor signaling at glutamatergic synapses. Correlated confocal and stochastic optical reconstruction microscopy (STORM) imaging did not reveal differences in the overall abundance of CB_1_ receptor levels at glutamatergic (i.e., VGluT1-positive) afferents of VTA dopamine neurons (Fig. 2B,C). Conversely, we found a substantial decrease in the nanoscale density of bassoon at VGluT1-positive axon terminals obtained from the PCE group (Fig. 2D,E), which is consistent with weaker glutamatergic synapses impinging on dopamine cells (24). Collectively, these data suggest that developmental remodeling of the presynaptic effector apparatus together with a diminished CB_1_ receptor signaling function might account for the absence of endocannabinoid-mediated DSE at glutamatergic afferents on dopamine neurons in PCE male VTA.

**Figure 2.**
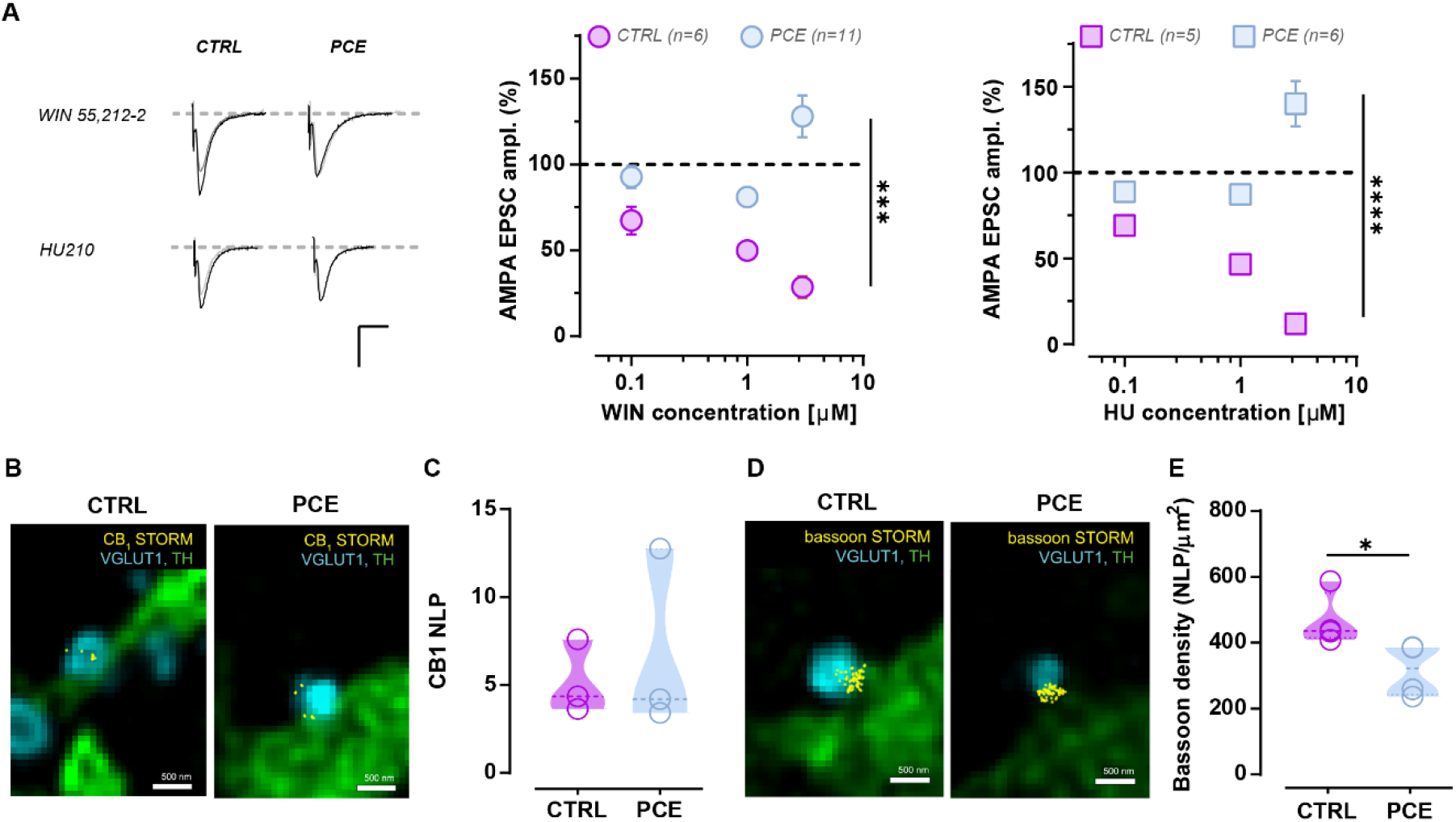
Prenatal THC pertubs cannabinoid tone at excitatory terminals on male dopaminergic neurons. A) AMPA EPSC traces (*left*) recorded before (black) and after (gray) bath application of either WIN 55,212-2 (WIN) or HU210 (HU) (both at 1 μM). Calibration bar, 5ms, 100pA. Concentration-response relationship for percentage decrease in AMPA EPSCs size produced by WIN 55,212-2 (*middle*) and HU210 (*right*). Each point shows the mean ± SEM of responses of different dopamine neurons from PCE and CTRL offspring (Effect of WIN 55,212-2 × treatment, *F*_(2,45)_=10,55, *P* = 0.0002; Effect of HU210 × treatment, *F*_(3,27)_=63,57, *P* < 0.0001 via two-way ANOVA). AMPA EPSC amplitude was normalized to the averaged value (dotted line) before drug application. Number in brackets indicates the number of cells. B) Correlated confocal and STORM super-resolution images of vGluT1-containing terminals (cyan) impinging on TH^+^ cells (green) in the VTA of CTRL and PCE animals. Yellow dots indicate CB1 localization points. C) Number of localization points (NLP) of the CB1 receptor staining in VGLUT1+ terminals (*N*=3 animals per group). D) Confocal image of VGLUT1 axon terminals (cyan) decorated with bassoon-STORM localization points (LPs) (yellow) marking the bouton active zone on TH+ cells (green) in the VTA of CTRL and PCE animals. E) Bassoon density in VGlut1^+^ active zones (two-sided Mann-Whitney *U-*test: P = 0.028; *N* = 3 animals per group). Unless otherwise indicated, graphs show violin plots (including the median values, and lower and upper quartiles) with each circle representing a single value. *\*P* < 0.05; *\*\*\*P* < 0.001; *\*\*\*\*P* < 0.0001.

### 3.2 Prenatal THC exposure switches retrograde signaling pathways for plasticity at excitatory terminals

To identify the molecular determinant of the endocannabinoid-independent LTD replacing the endocannabinoid-dependent DSE, we hypothesized that dopamine cells could recruit a different retrograde signal in response to a compromised CB_1_ signaling at glutamatergic afferents. One potential candidate could be nitric oxide (NO), which is released by VTA dopamine cells (31, 32). To test this hypothesis, we bath applied the non-selective NO synthase (NOS) inhibitor Nω-nitro-L-arginine methyl ester (L-NAME, 200 mM), which did not affect the endocannabinoid-dependent DSE in CTRL dopamine cells (Fig.3A). In contrast, L-NAME completely abolished the effects of postsynaptic depolarization on AMPA EPSCs recorded from PCE dopamine cells (Fig.3A). To identify the isoform of the NOS enzyme involved, we intracellularly applied the selective inhibitor of neuronal NOS (nNOS), Nω-Propyl-L-arginine (NPLA, 100 mM), through the patch pipette. NPLA blocked the effects on AMPA EPSCs in PCE dopamine cells without affecting DSE in CTRL group (Fig. 3B). Soluble guanylate cyclase (sGC), the enzyme responsible for cGMP synthesis, is an important molecular target of NO (31, 32). To assess whether cGMP plays a role in the novel endocannabinoid-independent NO-dependent plasticity, VTA slices were perfused with the sGC inhibitor 1H-[1,2,4]oxadiazolo[4,3-a]quinoxalin-1-one (ODQ; 1mM, 10 min preincubation). In the presence of ODQ, CTRL dopamine cells express intact DSE (~31%, Fig. 3C) with a similar magnitude to the one observed in the absence of ODQ (Fig. 1C) and in naïve animals (28, 30). Conversely, in the presence of ODQ, postsynaptic depolarization failed to change AMPA EPSCs recorded from PCE dopamine cells (Fig. 3C).

**Figure 3.**
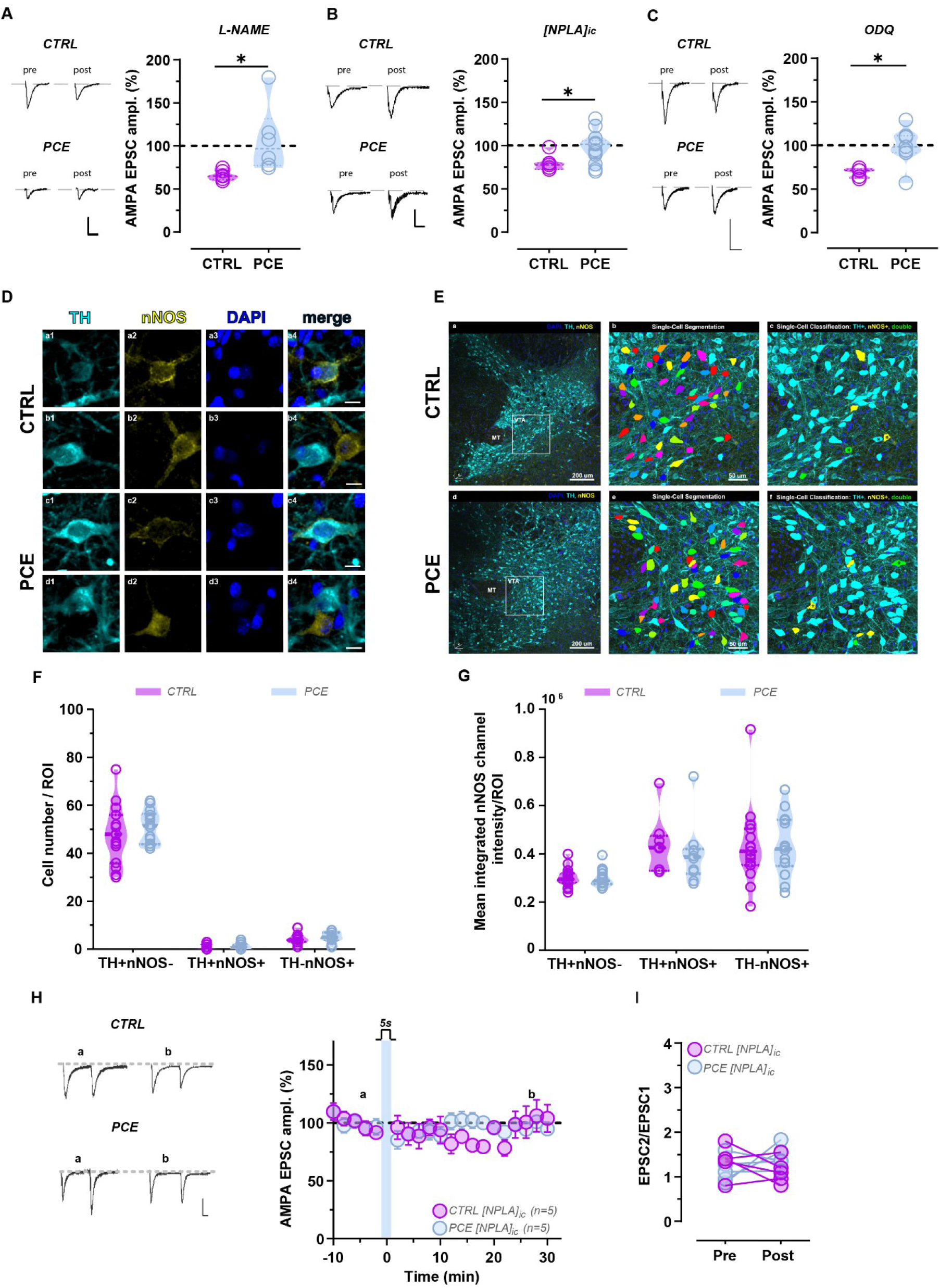
Prenatal THC switches retrograde signaling pathways for plasticity at excitatory terminals. A) Bath application of the NOS inhibitor L-NAME (200 mM) prevents DSE expression in male PCE dopamine neurons (Unpaired *t-*test: t=2.527, df=10, *P* = 0.03; *N* = 6 per group). *Left*, representative traces of AMPA EPSCs recorded before (pre) and after 5s depolarization during. Calibration bar, 10ms, 100pA. B) Intracellular inhibition of nNOS by NPLA (NPLAic, 100 µM) prevents DSE in PCE male dopamine cells (Unpaired *t*-test: t=2.546, df=18, *P* = 0.02; *N_CTRL_* = 7; *N_PCE_* = 13). *Left*, representative traces of AMPA EPSCs recorded before (pre) and after 5s depolarization. Calibration bar, 10ms, 100pA. C) Bath application of the sGC inhibitor ODQ (1 µM) prevents DSE expression in male PCE dopamine neurons (Unpaired *t*-test: t=2.912, df=10, *P* = 0.015; *N_CTRL_* = 5; *N_PCE_* = 7). *Left*, representative traces of AMPA EPSCs recorded before (pre) and after 5s depolarization. Calibration bar, 10 ms, 100 pA. D) Confocal images of immunofluorescent staining of TH+ (cyan, a1, b1, c1, d1), nNOS (yellow, a2, b2, c2, d2), DAPI (blue, a3, b3, c3, d3) and overlaid (a4, b4, c4, d4) channels from horizontal brain sections from CTRL (a-b) and PCE (c-d) animals. Calibration bar: 10 μm. E) Confocal images of immunofluorescent staining of TH (cyan) and nNOS (yellow) from horizontal brain sections of CTRL (a-c) and PCE (d-f) rats. b, e) Enlarged images from the VTA (the boxed area in a and c) are overlaid with deep-learning-based single-cell segmentation masks and c, f) machine-learning classification of TH^+^ (cyan) and nNOS^+^ (yellow) and double positive (green) cells. Asterisks indicate selected cells shown in panel D. F,G) Quantification of single-cell based segmentation and machine learning modes with three classes: TH+nNOS+, TH+nNOS− and TH−nNOS+ show that (F) there are no differences in the number of different class members(Pairwise Mann-Whitney *U-*test: TH+nNOS−, *P* = 0.406; TH+nNOS+, *P* = 0.453; TH−nNOS+, *P* = 0.537; *N_CTRL_* = 15; *N_PCE_* = 14) and (G) in the nNOS integrated intensity between CTRL and PCE (Pairwise Mann-Whitney *U-*test: TH+nNOS−, *P* = 0.711; TH+nNOS+, *P* = 0.408; TH−nNOS+, *P* = 0.810; *N_CTRL_* = 8-15; *N_PCE_* = 10-14). H) *Left*, traces show evoked paired AMPAR EPSCs obtained before (a) and after 30 min (b) of 5s depolarization in the presence of NPLAic. Calibration bar, 10ms, 100pA. *Right,* in PCE male dopamine cells, the NPLAic (100 µM) abolishes the induction of 5s depolarization induces a long-term depression of AMPA EPSCs. Each point represents the average of the mean EPSCs for 2 min-bin (±SEM) obtained from different cells. Number in brackets indicates the number of cells. I) The graph plots the paired-pulse ratio for each of the experiments in H before (pre) and after DSE (post) in male dopamine neurons. Unless otherwise indicated, graphs show violin plots (including the median values, and lower and upper quartiles) with each circle representing a single value. *\*P* < 0.05.

The expression of nNOS in VTA dopamine cells remains controversial (33, 34). While immunolabeling studies reported high expression of nNOS in GABAergic interneurons in the VTA (34), scRNAseq data shows *nos1-*expression in a select population of VTA neurons with moderate *th* (tyrosine hydroxylase, TH; the rate limiting enzyme in dopamine synthesis) mRNA expression (35). To address this issue at the protein level, we performed a confocal microscopy analysis of cells expressing nNOS and TH, and confirmed the colocalization between TH and nNOS proteins in a select population of cells in male rat VTA (Fig. 3D-F). Next, we examined the degree of colocalization by implementing a machine learning model with three classes (i.e., TH+nNOS+, TH+nNOS−, TH−nNOS+). We found that the integrated intensity of nNOS−immunolabeling was unaffected by PCE (Fig. 3E, G). Interestingly, while TH intensity did not change as a function of PCE TH+nNOS−cells (Fig. S2A), TH intensity level was found to be higher in PCE TH+nNOS+ cells (Fig. S2B). Next, intracellular nNOS inhibition with NPLA prevented the expression of the endocannabinoid-independent NO-dependent LTD in PCE dopamine cells, without modifying AMPA EPSC amplitude in CTRL dopamine cells (Fig. 3H,I). Altogether, these physiological and anatomical observations indicate that NO/cGMP signaling is required for LTD induction in PCE dopamine cells, although PCE does not change nNOS protein abundance.

To provide an independent evidence that dopamine neurons can produce NO, we used the cell-permeable fluorescent NO sensor, 4-Amino-5-Methylamino-2’,7’-Difluorofluorescein (DAF-FM) diacetate (10 mM, 1 hr preincubation) (33). We directly probed if significant NO concentrations are released upon postsynaptic depolarization of dopamine neurons in acute VTA slices (Fig 4A). Measurement of changes in fluorescence intensity revealed no changes in NO levels in CTRL VTA slices (Fig. 4B-D). In striking contrast, the same induction protocol produced a robust increase in NO concentrations in PCE VTA slices (Fig. 4B-D). Collectively, these results reveal that PCE fundamentally changes the rules for depolarization-induced synaptic plasticity in male dopamine neurons converting an endocannabinoid-mediated, NO-independent, DSE into an NO-mediated, endocannabinoid-independent, LTD.

**Figure 4.**
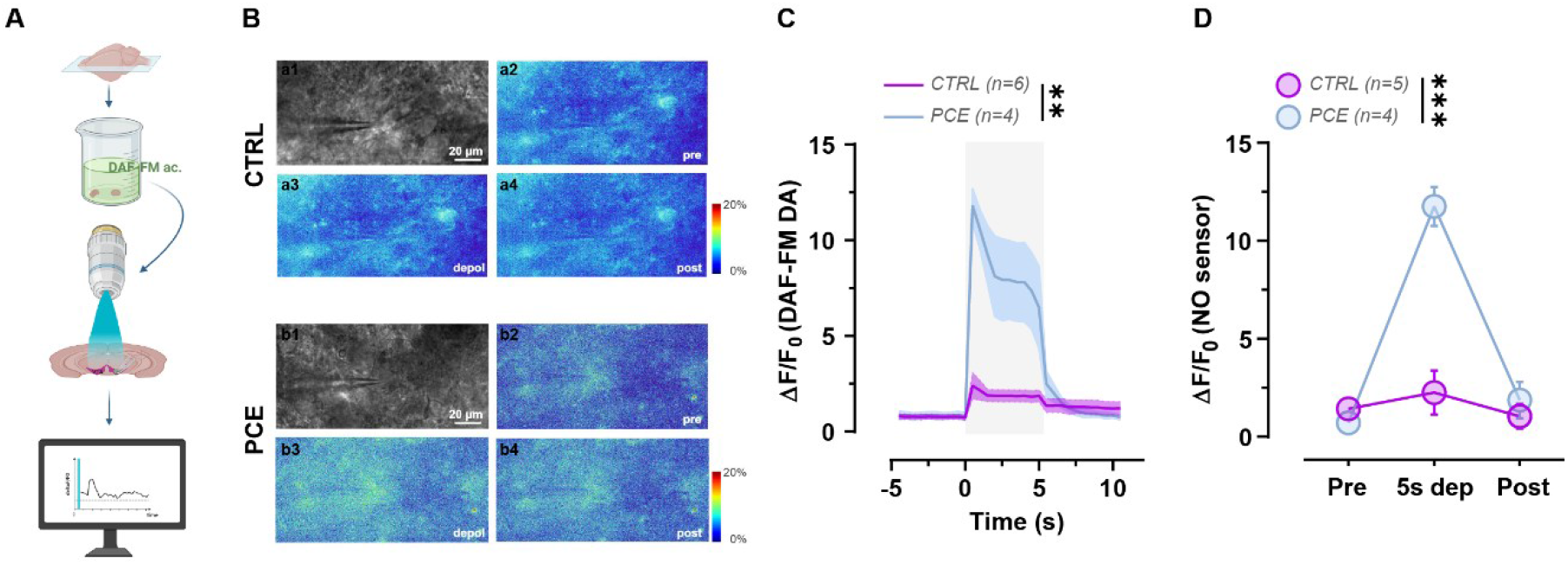
Prenatal THC increases depolarization-induced NO signal in VTA DA neurons. A) Schematic representation of the experimental protocol using the nitric oxide (NO)-sensitive probe DAF-FM diacetate on VTA midbrain sections. B) Representative grayscale images (a1, b1) and pseudocolor ΔF/F₀ maps from CTRL (a) and PCE (b) slices at baseline (pre; a2, b2), during 5-s depolarization (depol; a3, b3), and post-depolarization (post; a4, b4). C) Time course of stimulus-evoked NO production expressed as ΔF/F₀ over time in CTRL (pink) and PCE (blue) groups. Shaded area indicates the 5s depolarization period (Two-way RM ANOVA: treatment’s effect: F_(1,8)_ = 21.76, *P* = 0.0016; time’s effect: F_(1.378, 11.02)_ = 24.45, *P* = 0.0002; interaction between factors: F_(29,232)_ = 14.89, *P <* 0.0001; *N_CTRL_* = 6; *N_PCE_* = 4). C) Quantification of NO signals extracted from the time-lapse recordings shown in C, expressed as ΔF/F₀ during baseline (pre), depolarization (5s dep), and post-depolarization (post) periods. PCE slices display a significantly enhanced depolarization-evoked NO signal compared with CTRL, with no differences at baseline or post-stimulation (Two-way RM ANOVA: treatment’s effect: F_(1,7)_ = 33.57, *P* = 0.0007; time’s effect: F_(1.451, 10.162)_ = 44.76, *P <* 0.0001; interaction between factors: F_(2,14)_ = 30.74, *P <* 0.0001; *N_CTRL_* = 5; *N_PCE_* = 4). *\*\*\*P* < 0.001; *\*\*\*\*P* < 0.0001.

### 3.4 In utero THC induces a GABAb receptor-dependent form of hetero-synaptic plasticity in VTA dopamine cells

In the VTA, NO triggers LTP of inhibitory synapses (iLTP) (31, 32, 36–38). Having established that PCE dopamine cells substantially release NO in an activity-related (i.e. depolarization-dependent) manner to trigger LTD of excitatory inputs, we hypothesized that NO may also affect synaptic plasticity of inhibitory afferents. We, therefore, tested if the same depolarizing protocol may alter GABAergic transmission. Notably, we found that GABAa IPSCs undergo a form of LTP (~30%) in PCE, but not in CTRL dopamine cells (Fig. 5A). This iLTP was accompanied by a change of PPR toward depression (Fig.5B). Consistent with our previous results (Fig. 3), nNOS inhibition with intracellular NPLA application in PCE dopamine cells prevented the induction of iLTP (Fig. 5C-F). Next, we hypothesized that increased GABA release during iLTP could spill over to neighboring synapses to activate GABAb receptors, which would presynaptically reduce glutamate release (39) on PCE dopamine neurons. Importantly, in the presence of the GABAb receptor antagonist CGP35348 (100 mM), postsynaptic depolarization was ineffective on modifying AMPA EPSC amplitude (Fig. 5G) and PPR (Fig. 5I) in PCE male dopamine cells. Collectively, these results indicate that PCE induces a striking polarized switch in the direction of synaptic plasticity in dopaminergic neurons by inducing an NO-mediated LTP at inhibitory synapses and a hetero-synaptic GABAb-mediated LTD at excitatory synapses (Fig. S3).

**Figure 5.**
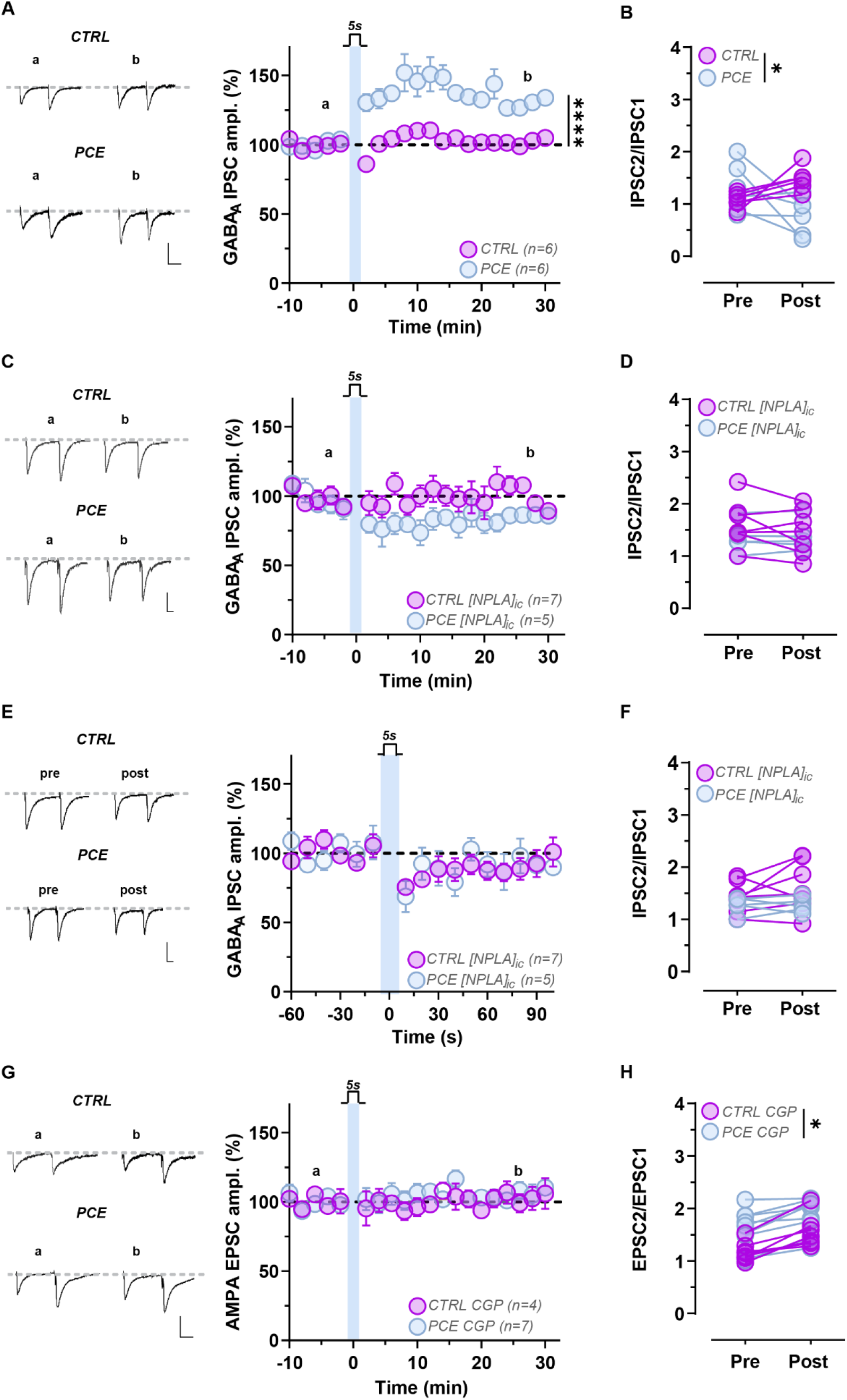
Prenatal THC induces synapse-specific and -dimorphic forms of plasticity in response to post-synaptic depolarization of dopamine neurons. A) *Left*, representative traces of paired GABAA IPSCs recorded pre (a) and 30 min post (b) depolarization from male CTRL and PCE dopamine neurons. Calibration bar, 10 ms, 100 pA. *Right*, time course of the effects of 5s depolarization on GABAA IPSC amplitude recorded from male dopamine neurons. Two-way mixed ANOVA: treatment’s effect: F_(1,10)_ = 37.21, *P* = 0.0001; time’s effect: F_(3.210,30.41)_ = 12.55, *P <* 0.0001; interaction between factors: F_(19,180)_ = 7.99, *P <* 0.0001; *N_CTRL_* = 5; *N* = 6 per group). GABAA IPSC amplitude was normalized to the averaged value (dotted line) before depolarization. B) Graphs plot the paired-pulse ratio (IPSC2/IPSC1) for each of the experiments in B before (pre) and 30 min after (post) depolarization (Two-way ANOVA: pre vs post’s effect: *F*_(1,20)_=0.20, *P* = 0.65; treatment’s effect: *F*_(1,20)_=3.22, *P* = 0.087; interaction between factors: *F*_(1,20)_=12.57, *P* = 0.002; Sidak’s multiple comparison test: post: *P* = 0.0024; *N* =6 per group). C) Effects of nNOS inhibition with NPLAic (100 µM) on GABAA IPSCs recorded in male dopamine neurons. *Left*, representative traces of paired GABAA IPSCs recorded pre and 30 min post depolarization from male dopamine neurons in the presence of NPLAic. Calibration bar, 10ms, 100pA. *Right*, time course of the effects of 5s depolarization on GABAA IPSC amplitude recorded from male dopamine neurons in the presence of NPLAic. IPSC amplitude was normalized to the averaged value (dotted line) before depolarization (Two-way ANOVA: treatment’s effect: *F*_(1,160)_=1.17, *P* = 0.28; time’s effect: *F*_(19,160)_=1.19, *P* = 0.27; interaction between factors: *F*_(19,160)_=0.91, *P* = 0.56; *N_CTRL_* = 7; *N_PCE_* = 5). D) Graph plots the paired-pulse ratio for each of the experiments shown before (pre) and 30 min after (post) depolarization in the presence of NPLAic (Two-way ANOVA: pre vs post’s effect: *F*_(1,8)_=0.05, *P* = 0.83; treatment’s effect: *F*_(1,8)_=0.26, *P* = 0.62; interaction between factors: *F*_(1,8)_=3.47, *P* = 0.099). E) Time course of DSI in male dopamine cells in the presence of NPLAic (two-way ANOVA: treatment’s effect: *F*_(1,208)_= 0.077, *P* = 0.78; time’s effect: *F*_(15,208)_= 2.217, *P* = 0.0069; interaction between factors: *F*_(15,208)_= 0.64, *P* = 0.83; *N_CTRL_* = 7; *N_PCE_* = 5). Each point represents the average of the mean IPSCs for the 10 sec-bin (±SEM) obtained from different cells. Number in brackets indicates the number of cells. *Left*, traces show evoked GABAA IPSCs obtained before (pre) and after (post) of 5s depolarization. Calibration bar, 10ms, 100pA. F) Graphs plot the paired-pulse ratio (IPSC2/IPSC1) for each of the experiments in E before (pre) and after (post) depolarization (Two-way ANOVA: pre vs post’s effect: *F*_(1,20)_=0.58, *P* = 0.45; treatment’s effect: *F*_(1,20)_=3.61, *P* = 0.071; interaction between factors: *F*_(1,20)_=0.48, *P* = 049).G) Effects of GABAB receptor antagonist CGP35348 (CGP, 100 mM) on AMPA EPSCs recorded in male dopamine neurons. *Left*, representative traces of paired EPSCs recorded pre (a) and 30 min post (b) depolarization from male dopamine neurons in the presence of CGP. Calibration bar, 10ms, 100pA. *Right*, time course of the effects of 5s depolarization on AMPA EPSC amplitude recorded from male dopamine neurons in the presence of CGP (Two-way ANOVA: treatment’s effect: *F*_(1,180)_=2.58, *P* = 0.10; time’s effect: *F*_(19,180)_=0.64, *P* = 0.87; interaction between factors: *F*_(19,180)_=0.43, *P* = 0.98). EPSC amplitude was normalized to the averaged value (dotted line) before depolarization. H) Graph plots the paired-pulse ratio for each of the experiments shown before (pre) and 30 min after (post) depolarization in the presence of CGP (Two-way ANOVA: pre vs post’s effect: *F*_(1,28)_=8.27, *P* = 0.007; treatment’s effect: *F*_(1,28)_=10.59, *P* = 0.003; interaction between factors: *F*_(1,28)_=0.21, *P* = 0.65; Sidak’s multiple comparison test: *P* = 0.029; *N_CTRL_* = 7; *N_PCE_* = 9). Unless otherwise indicated, each point of the time course represents the average of the mean EPSCs for the 2 min-bin (±SEM) obtained from different cells. Number in brackets indicates the number of cells. *\*P* < 0.05; *\*\*\*\*P* < 0.0001.

## 4. Discussion

Here we report that in utero THC triggers a fundamental reconfiguration of retrograde signaling mechanisms at synapses in the VTA of male preadolescent offspring that may coordinate heterosynaptic metaplasticity within the VTA circuitry when cannabinoid signaling is compromised. Such synapse-dimorphic and weight-dependent changes manifest upon depolarization of dopamine neurons and critically require postsynaptic activation of nNOS, NO release and cGMP synthesis. The observed form of iLTP mediates lateral inhibition of excitatory afferents as GABA acting on GABAb receptors accounts for this novel form of heterosynaptic LTD. These findings expand on the mechanisms accounting for the excitatory-to-inhibitory (E/I) imbalance of PCE VTA dopaminergic neurons (24) that are likely to contribute to male-specific behavioral consequences of maternal cannabis use. By disrupting the molecular logic of retrograde synaptic signaling within the male VTA, PCE heightens E/I ratio and susceptibility to stress, which impairs sensorimotor gating functions exclusively of the male progeny (25, 26).

Our data supports and extends the evidence that prenatal exposure to THC produces persistent alterations in nanoscale release machinery without affecting CB_1_ receptor levels (24, 40, 41). Since CB_1_ lowers initial release probability, it is plausible that in male VTA slices of PCE rats, which already display a weakened vGlut1 synaptic connectivity onto dopamine neurons (24), the dynamic range of action of CB_1_ receptors might be constrained to prevent the further lowering of glutamate release (23 and present data). Alternatively, an enhanced endocannabinoid tone at these synapses might occlude the effect of subsequent bath application of CB_1_ receptor agonists or of postsynaptic depolarization, thus providing an underlying mechanism for the observed persistent facilitation of these synapses. In this framework, PCE might interfere with the developing endocannabinoid system to produce at least two interconnected functional states where prior and tonic engagement of cannabinoid signaling may either trigger metaplasticity or represent a means for metaplastic control of endocannabinoid system itself (42). Hence, PCE synapse-dimorphic changes in CB_1_ receptor signaling occurring in the male VTA may play a role in dopamine cell disinhibition, by increasing and dampening cannabinoid signaling at inhibitory (24) and excitatory afferents (present data), respectively. Such a synapse-specific shift in the balance of cannabinoid control within the VTA circuitry might also provide a mechanism explaining why ineffective concentrations (and doses) of THC excite PCE VTA dopamine neurons both *ex vivo* and *in vivo*, and deteriorate sensorimotor gating functions in a male-specific manner (24, 25, 27).

The novel form of LTD here described, only exhibited by PCE male dopamine cells, might be part of those metaplastic mechanisms occurring at this synapse where low-frequency afferent stimulation cannot induce an endocannabinoid-dependent LTD but rather promotes LTP (24). Thus, this NO/cGMP-dependent and GABAb-mediated LTD could replace endocannabinoid-mediated LTD to recalibrate excitatory synaptic strength on PCE dopamine cells. Of note, a different rule applies to dopamine cells (naïve and CTRL), in which endocannabinoids mediate both a CB_1_-dependent DSE (present data and (28)) and a CB_1_-mediated LTD upon LFS (present data and (43)). Accordingly, nNOS inhibition does not affect basal GABA and AMPA postsynaptic currents and does not dampen CB_1_-mediated effects of postsynaptic depolarization (Fig. 2 and 5), which *per se* does not trigger NO release. Thus, constitutive nNOS activity is not necessary to maintain basal levels of presynaptic GABA and glutamate release on VTA dopamine neurons. The specific post-translational or cell physiological mechanism underlying how PCE upregulates nNOS function in an activity-dependent manner remains to be determined. Nonetheless, our real-time imaging data demonstrate a striking NO spike triggered in male PCE VTA dopamine neurons. Such high levels of NO generation are suggestive of abnormal intracellular calcium levels in PCE male dopamine neurons, consistent with the expression of immature forms of ionotropic glutamatergic receptors and a leftward shift in the threshold for plasticity induction (24). Since THC can influence mitochondrial function (44), one intriguing possibility would be that PCE deregulate mitochondrial dynamic control of intracellular calcium homeostasis. While a deeper understanding of PCE-induced forms of metaplasticity deserves further elucidation, the observation that NO can be generated under conditions that usually trigger endocannabinoid release (28, 30, 45–48) may suggest that PCE biases retrograde signaling pathways to match changes in presynaptic CB_1_ receptor control efficacy. One may, therefore, speculate that PCE allows for a functional switch in the identity of retrograde signaling molecules (NO vs endocannabinoids) released upon depolarization of dopamine cells. Behaviorally, this would oppose the effects of acute stress on both excitatory (49, 50) and inhibitory (38) inputs on male dopamine cells in PCE individuals where the range of plasticity is constrained (24), thus contributing to stress-related phenotypes (25, 26).

Here, we extend previous observations that an NO-dependent signaling pathway can potentiate GABA synapses (i.e., iLTP) on VTA dopamine neurons (31, 36, 51). This novel NO-dependent presynaptic increase of GABA release, a phenomenon that could only be observed in PCE male dopamine neurons, likely explains the expression of LTD, which requires presynaptic GABAb-receptor activation. In this scenario, LTD could be envisaged as a form of lateral inhibition to regulate dopamine neuron activity when the presynaptic gain of cannabinoid signaling is compromised. This plasticity requires NO produced by dopamine cells, which can easily diffuse to activate sGC generating cGMP and to promote GABA release. Whether this is the precise sequence of signaling events remains to be disclosed. Nonetheless, it is noteworthy that potentiation of inhibitory afferents on dopamine cells operated by NO/cGMP signaling is a compelling example of synapse-specificity within the VTA: in fact, only nucleus accumbens and VTA GABAa synapses on dopamine cells can be potentiated by this signaling cascade (32, 37, 52). This suggests that such a shared plasticity mechanism accounting for the observed lateral inhibition (i.e., LTD) not only is spatially segregated based on dopamine cell location (i.e., lateral VTA), but it may also depend on their incoming inputs (37, 52). Importantly, whether such forms of metaplasticity are driven by changes in endocannabinoid signaling triggered by or resulting from PCE remains to be elucidated. Nonetheless, given that PCE modifies VTA circuit architecture (24), it is plausible that it also shapes its modularity in an input/output-specific manner to constrain the dynamic range of synaptic activity within the VTA.

In conclusion, these findings gain further insights into the molecular and synaptic mechanisms that are induced by PCE on dopamine neuronal trajectories. Expanding the evidence on the lifelong perturbations determined by the main psychoactive ingredient of cannabis is important considering that cannabis use among pregnant women is increasing along with its social acceptability and accessibility (53–56). Since children affected by in utero THC exposure are known to be predisposed to adverse neuropsychiatric outcomes (11–13, 15–20, 57), our study paves the way for further investigations to uncover how PCE affects neurodevelopmental trajectories and contribute to increased susceptibility to psychiatric disorders.

## Author Contributions

V.S., F.B. and V.M conducted experiments and analyzed data. M.M. designed the study, performed some of the experiments, analyzed the data, wrote the manuscript and supervised the project. I.K. provided the expertise for STORM and confocal experiments and contributed to manuscript preparation.

## Author approvals

This manuscript has not been accepted or published elsewhere. All authors have read and approved the final manuscript.

## Acknowledgments

This study was supported by the Horizon Europe 2022 Excellent Science - European Research Council (101088207 to M.M.), the National Institutes of Health (P30DA056410) and by the NKFIH EXCELLENCE program (151377) to IK, who holds the Naus Family Chair in Addiction Sciences in the Department of Psychological and Brain Sciences at Indiana University Bloomington. MV received funding from NKFIH OTKA PD (147127) and from the Hungarian Academy of Sciences - Grant for Researchers Raising Small Children mechanism.

## Data availability statement

Data can be provided from the corresponding authors upon reasonable request.

## 5 Figures and tables

**Figure Supplementary 1.**
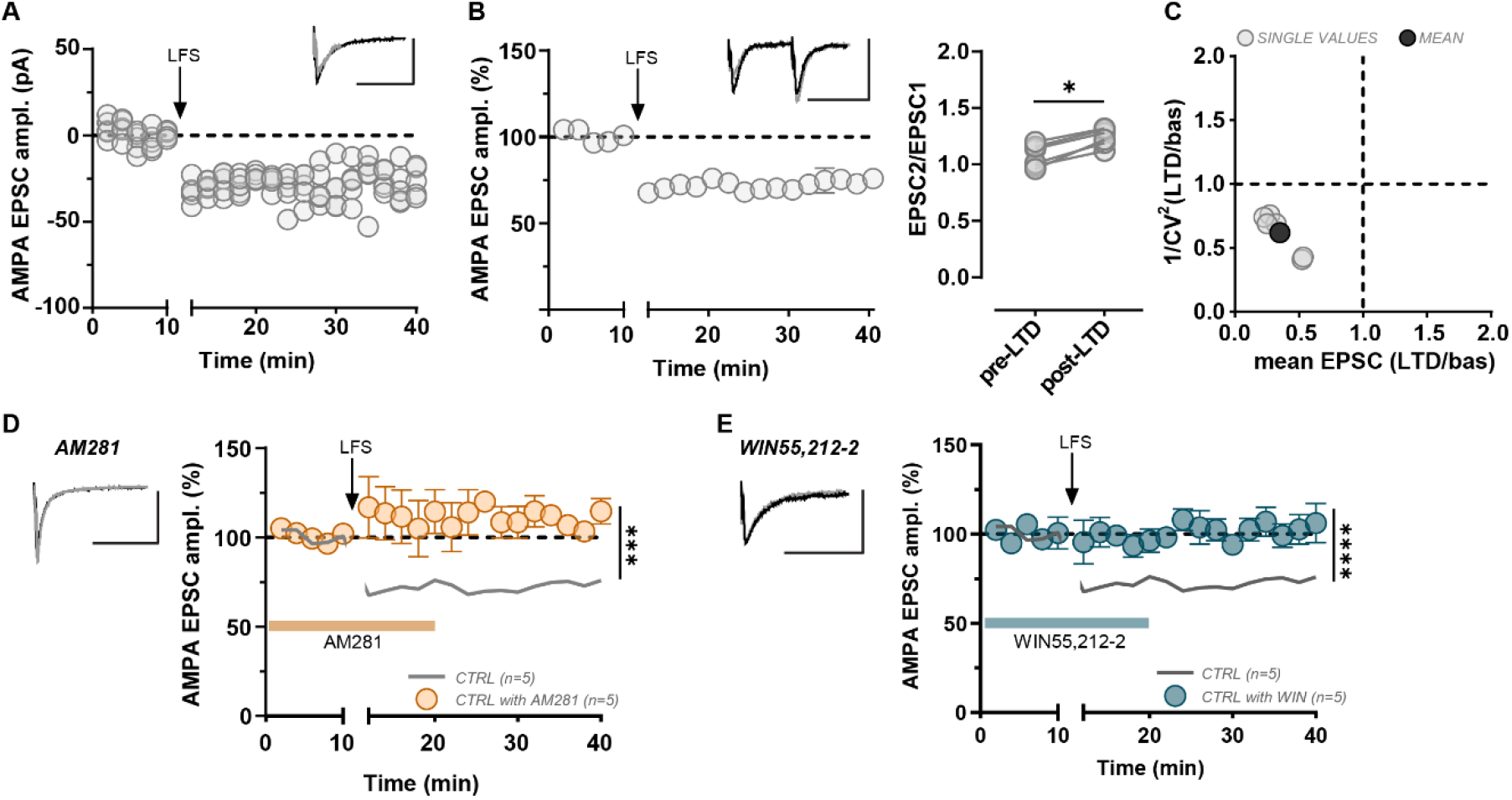
A) Effects of low-frequency afferent stimulation (LFS; 1 Hz at the arrow) on AMPA EPSC amplitude recorded in VTA DA neurons of naive animals (*N* = 5). Data represents the response of each cell recorded. Inset shows traces (−70 mV) from before (black) and after LFS (grey). Calibration bar, 50 ms, 100 pA. B) *Left*, LFS induced LTD on AMPA EPSC recorded from VTA DA neurons of naive preadolescent rats. Data represents the average ± s.e.m. of experiments displayed in A. Inset shows the traces of AMPA EPSCs (−70 mV) recorded before (black) and after LFS (grey). Calibration bar, 50 ms, 100 pA. *Right*, Graph plots the paired-pulse ratio (unpaired *t*-test: t=3.125, df=10, *P* = 0.01; N *= 6*) for each of the experiments shown before (pre-LTD) and 30 min after LFS (post-LTD). C) Graph representing changes in coefficient of variation (CV) of the experiments in A,B. 1/CV^2^ (LTD/bas) represents the ratio between 1/CV squared, obtained during LTD in response to LFS. Mean EPSC (LTD/bas) represents the ratio between mean EPSC amplitude, measured during LTD and before LFS. D) *Left*, representative traces of AMPA EPSC recorded before (black) and after LFS (grey) following bath application of AM281 (500nM). Calibration bar, 50 ms, 100 pA. *Right*, AM281 abolishes LFS-induced Ltd on AMPA EPSC recorded from VTA DA neurons (Two-way ANOVA: AM281’s effect: F_(1,160)_= 159.2, *P* < 0.0001; time’s effect F_(19,160)_= 0.85, *P* = 0.64; interaction between factors: F_(19,160)_= 2.93, *P* = 0.0001). E) *Left*, representative traces of AMPA EPSC recorded before (black) and after LFS (grey) in presence of WIN55,212-2 (1pM). Calibration bar, 50ms, 100pA. *Right*, WIN55,212-2 occludes LFS-induced LTD on AMPA EPSC recorded from VTA DA neurons (Two-way ANOVA: WIN55,212-2’s effect: F_(1,140)_= 164, *P* < 0.0001; time’s effect F_(19,140)_= 3.31, *P* < 0.0001; interaction between factors: F_(19,140)_= 3.53, *P* < 0.0001). Experiments were repeated independently with similar results obtained. Unless otherwise indicated, each point of the time course represents the average of the mean EPSCs for the 2 min-bin (±SEM) obtained from different cells. Number in brackets indicates the number of cells. *\*P* < 0.05; *\*\*\*P* < 0.001; *\*\*\*\*P* < 0.0001.

**Figure Supplementary 2.**
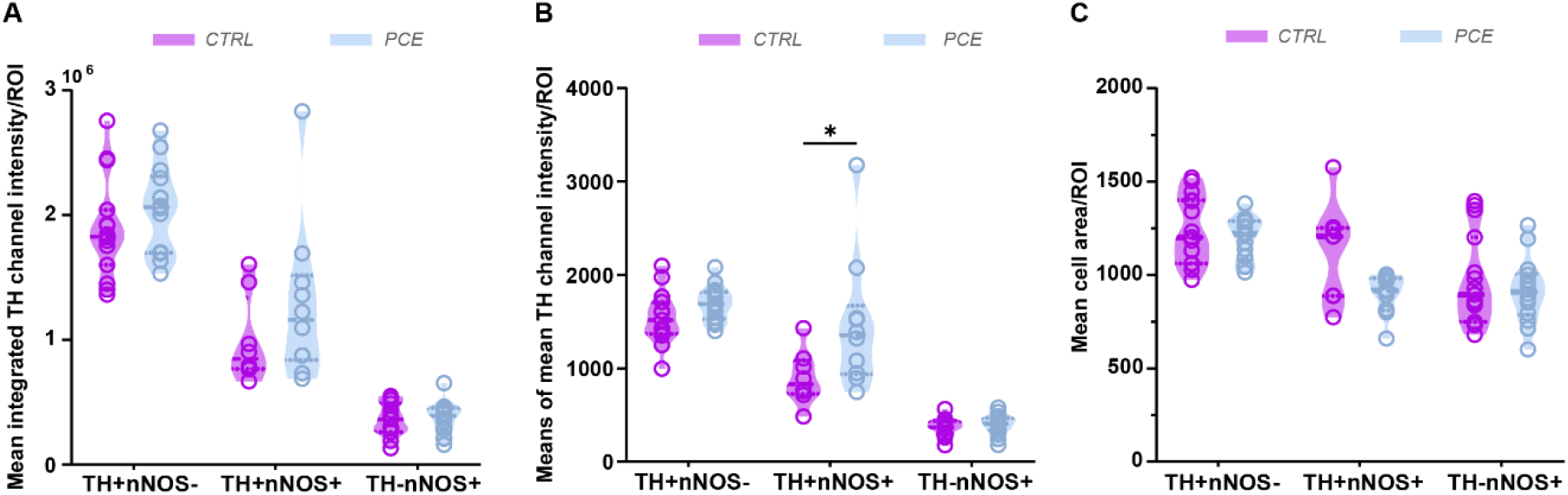
Quantification of single-cell based segmentation and machine learning modes with three classes: TH+nNOS− (*N_CTRL_*= 15; *N_PCE_* = 14), TH+nNOS+ (*N_CTRL_*= 8; *N_PCE_* = 10) and TH−nNOS+ (*N_CTRL_*= 15; *N_PCE_* = 14) show that (A) there are no differences in the TH intensity between CTRL and PCE (Pairwise Mann-Whitney *U-*test: TH+nNOS−, *P* = 0.285; TH+nNOS+, *P* = 0.408; TH−nNOS+, *P* = 0.844), whereas (B) the mean TH level is higher in the double positive PCE cells than CTRL (Pairwise Mann-Whitney *U-*test: TH+nNOS−, *P* = 0.085; TH+nNOS+, *P* = 0.034; TH−nNOS+, *P* = 0.395). Panel (C) shows no significant difference in cell area between CTRL and PCE treatments (Pairwise Mann–Whitney *U-*test: TH+nNOS−, *P* = 0.585; TH+nNOS+, *P* = 0.146; TH−nNOS+, *P* = 0.395). However, a non-significant trend toward smaller cell area is observed in the TH+nNOS+ PCE cells compared to CTRL, which may contribute to the significant difference observed in mean TH intensity in panel B. Graphs show violin plots (including the median values, and lower and upper quartiles) with each circle representing a single value. *\*P* < 0.05.

**Figure Supplementary 3.**
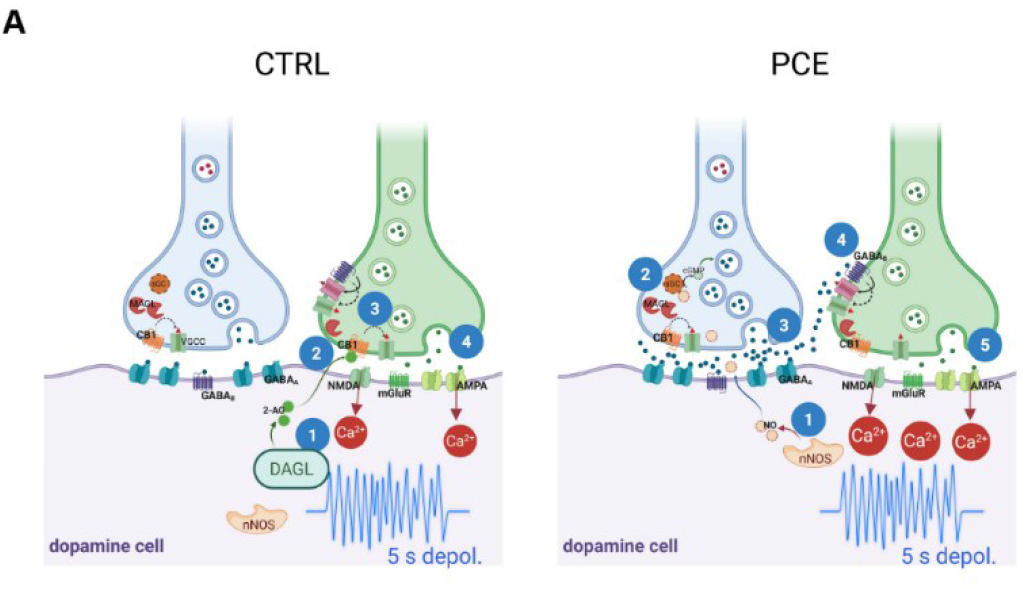
Proposed model for the switch in retrograde endocannabinoid-nitric oxide signaling in prenatally THC exposed (PCE) male rats. *Left,* At excitatory synapses on ventral tegmental area (VTA) dopamine neurons, 5s postsynaptic depolarization (5s depol.) leads to an endocannabinoid (i.e., 2-arachidonoylglycerol, 2-AG)-mediated form of short-term synaptic plasticity (i.e., depolarization induces-suppression of excitation (DSE)(28, 30). 5s-depol activates diacylglycerol lipase (DAGL) (1) and 2-AG migrates retrogradely to bind to presynaptic type-1 cannabinoid receptors (CB1) on glutamate terminals (3) to depress neurotransmitter release (4). *Right,* in PCE VTA DA cells, 5s-depol stimulates the activity of neuronal nitric oxide synthase (nNOS) thus releasing NO (1). Mobilized NO, in turn, activates the soluble guanylate cyclase (sGC) on GABA afferents (2) to induce long-term synaptic potentiation and presynaptic release of GABA (3) on DA neurons. GABA spillover binds to and activate GABAB receptors of neighboring excitatory synapses (4) to induce their long-term depression (5).

**Supplementary Table 1.**

| Target | Host species | Distributor | Cat.no. | Concentration |
| --- | --- | --- | --- | --- |
| **Primary antibodies** |  |  |  |  |
| Bassoon | Rabbit | Millipore | ABN255 | 1:1000 |
| Tyrosine Hydroxylase (TH) | Mouse | Immunostar | 22941 | 1:5000 |
| Vesicular Glutamate Transporter 1 (VGLUT1) | Guinea pig | Synaptic Systems | 135 304 | 1:5000 |
| Cannabinoid receptor type 1 | Rabbit | ImmunoGenes | - | 1:1000 |
| **Secondary antibodies** |  |  |  |  |
| Alexa647-conjugated anti-rabbit | Donkey | Jackson | 711-605-152 | 1:400 |
| Alexa488-conjugated anti-mouse | Donkey | Jackson | 715-545-150 | 1:400 |
| CF568-conjugated anti-guinea pig | Donkey | Biotium | 20377-50ul | 1:1000 |

